# cryoJAX: A Cryo-electron Microscopy Image Simulation Library In JAX

**DOI:** 10.1101/2025.10.23.682564

**Authors:** Michael J. O’Brien, David Silva-Sánchez, Geoffrey Woollard, Kwanghwi Je, Sonya M. Hanson, Daniel J. Needleman, Pilar Cossio, Erik Henning Thiede, Miro A. Astore

**Affiliations:** Department of Physics, Harvard University, Cambridge, 02143, MA, USA; Department of Applied Mathematics, Yale University, New Haven, 06520, CT, USA; Department of Computer Science, University of British Columbia, Vancouver, V6T 1Z4, British Columbia, Canada; Department of Chemistry and Chemical Biology, Cornell University, Ithaca, 14853, NY, USA; Center for Computational Biology, Flatiron Institute, New York, 10010, NY, USA; Department of Molecular and Cellular Biology, Harvard University, Cambridge, 02143, MA, USA; John A. Paulson School of Engineering and Applied Sciences, Harvard University, Cambridge, 02143, MA, USA; Center for Computational Mathematics, Flatiron Institute, New York, 10010, NY, USA

**Keywords:** cryo-EM, image simulation, automatic differentiation, software library, biophysics

## Abstract

While cryo-electron microscopy (cryo-EM) has come to prominence in the last decade due to its ability to resolve biomolecular complexes at atomic resolution, advancements in experimental and computational methods have made cryo-EM promising for investigating intracellular organization and heterogeneous molecular states. A primary challenge for these alternative applications is the development of techniques for cryo-EM data analysis, which are very computationally demanding. To this end, it is advantageous to leverage advanced scientific computing frameworks for statistical analysis. One such framework is JAX, an emerging array-oriented Python numerical computing package for automatic differentiation and vectorization with a growing ecosystem for statistical inference and machine learning. We have developed cryoJAX, a cryo-EM image simulation library for building computational data analysis applications in JAX. CryoJAX is a flexible modeling language for cryo-EM image formation and therefore can support a wide range of data analysis downstream. By integrating with the JAX ecosystem, cryoJAX enables the development and deployment of algorithms for the growing breadth of scientific applications for cryo-EM.

**Synopsis:** The authors have developed cryoJAX, a cryo-EM image simulation library for developing data analysis techniques across cryo-EM modalities. CryoJAX is built on JAX, an emerging scientific computing framework in Python well suited for cryo-EM data analysis.

## 1 Introduction

Reconstruction of high-resolution biomolecular structure via cryo-electron microscopy (cryo-EM) has revolutionized structural biology in the last decade. This transformation has been made possible by dramatic improvements in experimental and computational methods for cryo-EM. Now cryo-EM is becoming a promising technique for investigating a broad range of biophysical phenomena, including intracellular organization (Lucas *et al*., 2021) and heterogeneous molecular states (Tang *et al*., 2023*b*). One of the main challenges is properly analyzing cryo-EM images, which come in large volumes and are high-resolution but low signal-to-noise.

Most existing software tools focus on single-particle reconstruction (Tang *et al*., 2007; Scheres *et al*., 2008; Scheres, 2012; Punjani *et al*., 2017*a*; Grant *et al*., 2018), but novel algorithms are unlocking new ways to use cryo-EM data. The single-molecule nature of cryo-EM data enables statistical inference of molecular properties, which requires models of cryo-EM image formation. Statistical inference built on cryo-EM image simulation is the backbone of recent applications of cryo-EM data analysis for emerging scientific applications (Lucas *et al*., 2021; Tang *et al*., 2023*b*). These applications are very computationally demanding. Therefore, it would be advantageous to integrate cryo-EM image simulation with advanced scientific computing frameworks for statistical analysis.

One such framework is JAX, a Python numerical computing library for just-in-time (JIT) compilation, automatic differentiation, vectorization, and parallelization across GPUs (Bradbury *et al*., 2018). These features support libraries for optimization (DeepMind *et al*., 2020; Rader *et al*., 2024), linear solvers (Rader *et al*., 2023), Monte-Carlo sampling (Phan *et al*., 2019; Cabezas *et al*., 2024), and machine learning (Heek *et al*., 2024; Kidger & Garcia, 2021). Other scientific disciplines have implemented physical modeling in JAX and used it to demonstrate the power of automatic differentiation for scientific applications (Brenner & King, 2023).

Methods for the reconstruction of continuous conformational heterogeneity from cryo-EM data have already demonstrated the power of combining image simulation with frameworks like JAX (Zhong *et al*., 2021; Li *et al*., 2024; Schwab *et al*., 2024; Gilles & Singer, 2025; Herreros *et al*., 2025*a*). The cryoDRGN software (Zhong *et al*., 2021) is built on PyTorch (Paszke *et al*., 2019), a Python-based framework with analogous features to JAX. To train its neural networks to represent 3D volumes, cryoDRGN uses automatic differentiation to compute gradients of a loss function with respect to neural network parameters. More generally, automatic differentiation makes it possible to use gradient-based statistical inference algorithms for cryo-EM data analysis of vast numbers of parameters. An image simulation library that integrates with these scientific computing resources could be used to build new analysis techniques, but current implementations are embedded within 3D reconstruction software and not designed for use across contexts.

We present cryoJAX, a flexible image simulation library in JAX. CryoJAX is a modeling framework for cryo-EM images and structures that is designed to be extended into external workflows and libraries that leverage JAX. It implements established image simulation models and algorithms, as well as a framework for implementing new models and algorithms. CryoJAX does not implement any particular data analysis, such as single-particle reconstruction. Rather, cryoJAX for modeling cryo-EM data and JAX for scientific computing is the foundation for cryo-EM data analysis. CryoJAX is positioned as a powerful tool for studying cryo-EM data for its emerging scientific applications.

## 2 Results

### 2.1 Overview of cryoJAX

CryoJAX is a cryo-EM image simulation library designed for building cryo-EM data analysis algorithms (Figure 1). A data analysis algorithm using cryoJAX takes cryo-EM images and prior information about them as input (Figure 1 A). It is advantageous to leverage existing frameworks for estimating information about the images. Information of the structures present in cryo-EM images can be derived from multiple sources, such as prior experiments, protein generative models (Raghu *et al*., 2025), or molecular dynamics simulations (Tang *et al*., 2023*a*). Information of imaging parameters can be estimated from existing cryo-EM software, such as for contrast transfer function (CTF) estimation (Rohou & Grigorieff, 2015).

**Figure 1.**
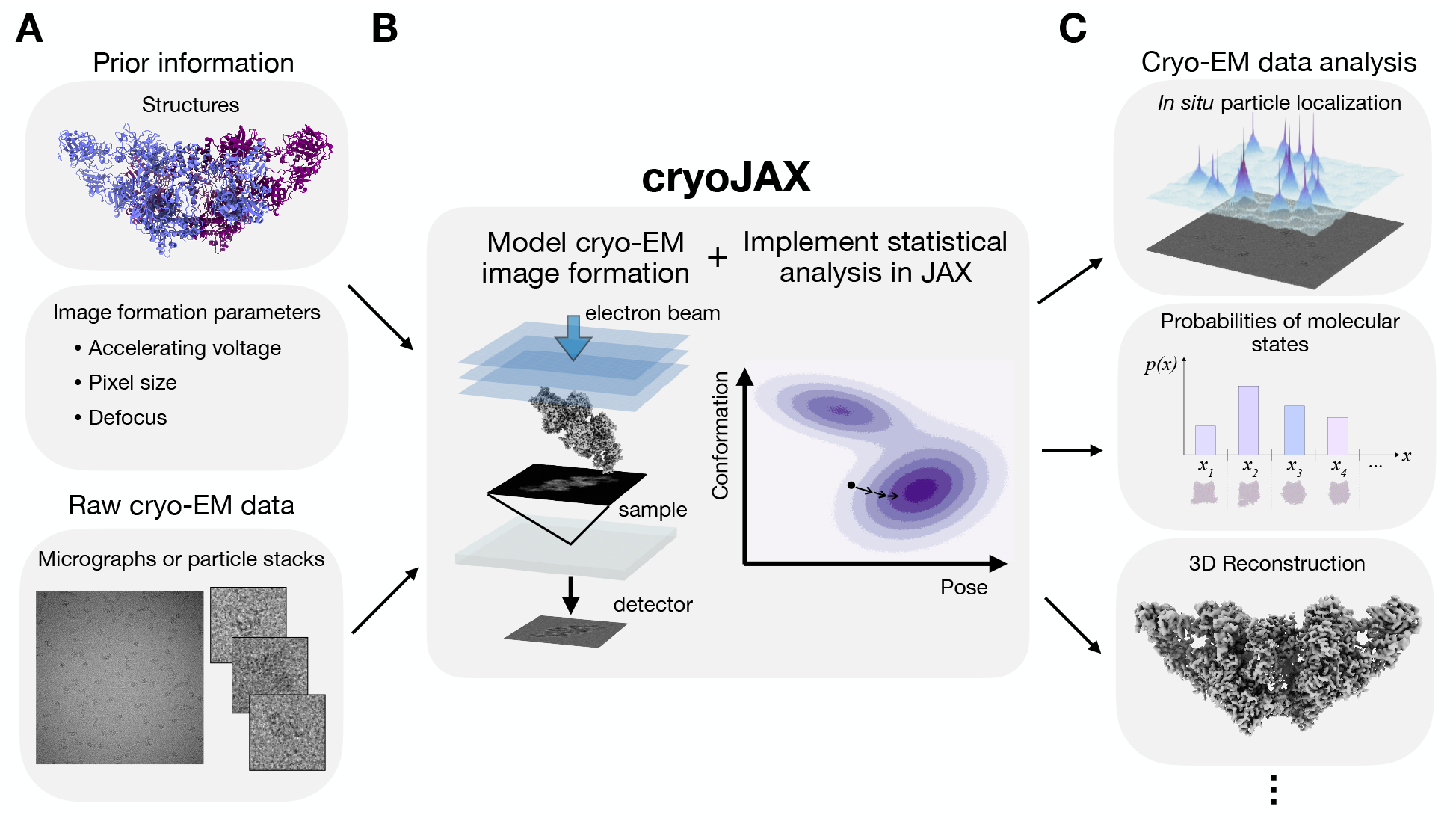
CryoJAX is an image simulation library for building cryo-EM data analysis. **A**. Building data analysis with cryoJAX starts by loading cryo-EM images and prior information about them into Python. **B**. Using these inputs, cryoJAX implements models for cryo-EM image formation and integrates with JAX for statistical analysis with high computational demands. This forms a foundation for cryo-EM data analysis applications. **C**. Using cryoJAX, it is possible to build a variety of cryo-EM data analysis techniques, including methods for *in situ* particle localization, inferring probabilities of molecular states, and 3D reconstruction.

Using these inputs, cryoJAX forms a foundation for cryo-EM data analysis applications (Figure 1 B). CryoJAX is an image simulation library that is highly flexible; it includes a range of well-established cryo-EM image simulation models and algorithms, as well as an interface for implementing custom methods external to the core library. The cryoJAX codebase is a framework for simulating single cryo-EM images, yet JAX makes it possible to build powerful and highly complex programs. JAX provides JIT compilation for C++/CUDA-level performance, automatic differentiation for gradient-based analysis of cryoJAX models in high-dimensional parameter spaces, and automatic vectorization for implementing complex workflows. With the ability to arbitrarily compose these transformations, JAX is well suited for the varied demands of cryo-EM data analysis.

CryoJAX is capable of supporting data analysis across applications of cryo-EM, such as for *in situ* particle localization (Lucas *et al*., 2021), inferring probabilities of molecular states (Tang *et al*., 2023*b*), or 3D reconstruction (Figure 1 C). These different applications are united by a need for models of cryo-EM image formation and advanced computational techniques, which is fulfilled by cryoJAX. For example, models of image formation facilitate Bayesian inference algorithms through a definition of the likelihood, and JAX is an emerging platform for Bayesian inference in the physical sciences. Indeed, JAX has already been leveraged in cryo-EM by the Bayesian heterogeneous reconstruction method implemented in RECOVAR (Gilles & Singer, 2025).

### 2.2 Image simulation features and implementation

The cryoJAX library implements models and algorithms for simulating cryo-EM images so that data analysis applications may be built using the scientific computing resources of JAX. Under an approximation of weak electron-specimen interaction, the formation of contrast in cryo-EM images may be approximated as a linear process: projection of a molecule’s electrostatic potential 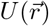 (Vulović *et al*., 2013) followed by convolution with the microscope’s point spread function, which in reciprocal space is the contrast transfer function (CTF). In many cases, the CTF can be expressed as sin *χ*, where *χ* = *χ*(*q*_*x*_, *q*_*y*_) is known as the aberration function (Rohou & Grigorieff, 2015) and (*q*_*x*_, *q*_*y*_) are reciprocal space coordinates. Considering a rotation **R** and translation ^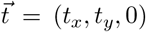^ of *U*, the contrast *C*(*x, y*) measured in lab-frame coordinates 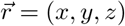 is

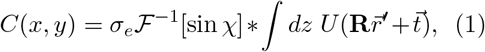

where 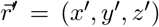 are body-frame coordinates, 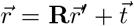 transforms from the body-frame to the lab-frame, *σ*_*e*_ is referred to as the electron interaction constant, and ℱ and ∗ are Fourier transform and convolution operations, respectively (Spence, 2013).

In practice, there are many ways to implement the volume *U*, the aberration function *χ*, and the rotation **R** (Figure 2 A), and the optimal choice depends on the details of a data analysis. To support data analysis with broad goals, cryoJAX is a modular and extensible framework for different image formation models and algorithms. For example, methods which study continuous conformational heterogeneity represent molecular volumes U in different ways (Donnat *et al*., 2022; Chen *et al*., 2023). Some use voxel maps (Gilles & Singer, 2025) or atomic coordinates (Vuillemot *et al*., 2023; Chen, 2025), but examples also exist which use deformation fields (Herreros *et al*., 2025*b*), neural networks (Zhong *et al*., 2021), and non-linear latent spaces (Ojha *et al*., 2025). Similarly, depending on the resolution of the data analysis, different models for the aberration function *χ* may be appropriate. At lower resolutions, it may suffice to only take into account defocus and spherical aberration, but at higher resolutions it can become necessary to model astigmatism and beam tilt. Finally, parametrizing the rotation **R** can be highly application-dependent. For example, cleverly parametrizing the space of rotations results in increased computational efficiency or improved geometry for inference (Barnett *et al*., 2017; Hanson & Hanson, 2022; Woollard *et al*., 2025*b*).

**Figure 2.**
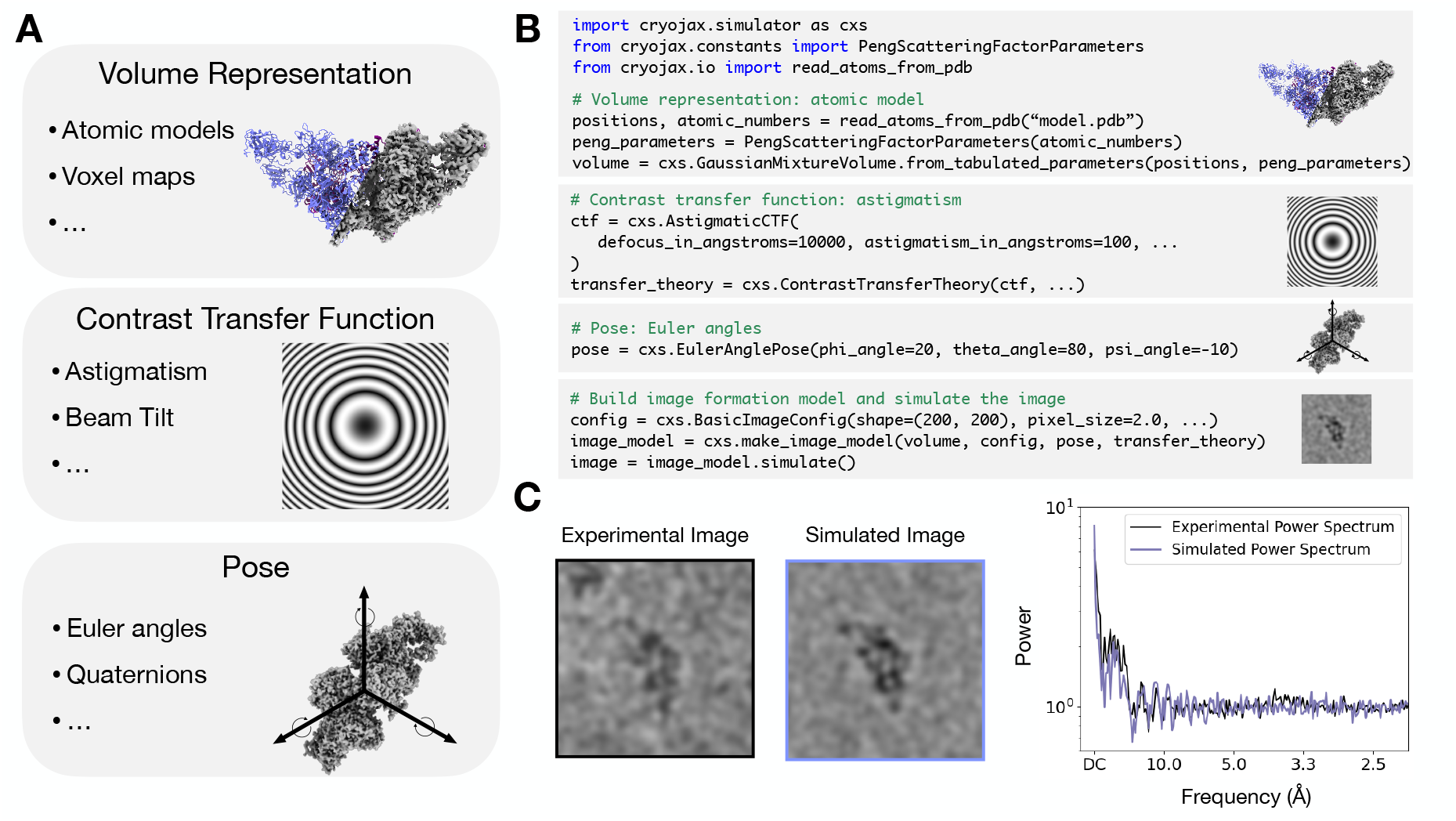
The cryoJAX library implements flexible but simple image simulation. **A.** In cryo-EM image simulation, there are multiple ways to parametrize volumes, contrast transfer functions (CTFs), and poses. Since the optimal choice varies based on the data analysis, cryoJAX is a framework for different implementations. **B**. One possible implementation of image simulation described in Equation 1 is easily invoked in a few lines of code by choosing a volume, CTF, and pose. **C**. The code from B. simulates images comparable to images in an experimental dataset of thyroglobulin (EMPIAR-10833). Power spectra are computed using a function in the cryoJAX image manipulation submodule, `cryojax.ndimage`. Images are displayed using a low-pass filter.

CryoJAX includes a core set of concrete implementations for each modeling component in Equation 1, as well as an API for writing new ones downstream via abstract base classes (Table 1). Abstract base classes define the interface that needs to be implemented to integrate with the cryoJAX image simulator. For example the `AbstractVolumeRepresentation` is the abstract base class for modeling volumes in cryoJAX. One concrete implementation of this in cryoJAX is the `GaussianMixtureVolume` class. This is an atomic model for the electrostatic potential approximated as the sum of Gaussian functions for each atom, which may be obtained from tabulated electron scattering factors (Appendix A). The current release of cryoJAX can compute projections of this volume analytically by computing the integral in Equation 1 or numerically by discretizing onto a 3D voxel grid, loading into a `FourierVoxelGridVolume`, and extracting Fourier slices.

**Table 1:** A summary of modeling components, abstract base classes, and selected concrete implementations in the cryoJAX library. Additional implementations can be incorporated in the future, and custom implementations may be created downstream to the cryoJAX core library.

| Modeling component | Abstract Base Class | Concrete Implementations |
| --- | --- | --- |
| Volume $U$ | `AbstractVolumeRepresentation` | • `FourierVoxelGridVolume`<br>• `GaussianMixtureVolume` |
| Pose $\mathbf{R}, \vec{t}$ | `AbstractPose` | • `EulerAnglePose`<br>• `QuaternionPose` |
| CTF $\sin \chi$ | `AbstractCTF` | • `AstigmaticCTF` |

**Table 2:** Compute times for simulating a single image with and without JIT compilation in cryoJAX.

| Method | CPU (ms) | GPU (ms) |
| --- | --- | --- |
| `GaussianMixtureVolume` (no JIT) | $216.2 \pm 10.8$ | $253.5 \pm 1.4$ |
| `GaussianMixtureVolume` (JIT) | $10.6 \pm 1.1$ | $0.58 \pm 0.09$ |
| `FourierVoxelGridVolume` (no JIT) | $302.4 \pm 25.4$ | $367.3 \pm 1.5$ |
| `FourierVoxelGridVolume` (JIT) | $5.53 \pm 0.27$ | $0.45 \pm 0.07$ |

Rotations may also be parametrized differently in cryoJAX through the `AbstractPose` interface. Euler angles are implemented in the `EulerAnglePose` class or quaternions in the `QuaternionPose`. Internally, each parametrization converts to a `cryojax.rotations.SO3` class, which rotates 3D vectors by multiplying quaternions and is based on the implementations in the JAX package `jaxlie` (Yi *et al*., 2021). The `AbstractPose` includes a method that inverts the rotation, as it may be necessary to do so to match projections from different volume representations.

The `AbstractCTF` class allows for modularity in CTF implementation via the aberration function *χ*. CryoJAX includes a core implementation `AstigmaticCTF`, which is implemented as in CTFFIND4 (Rohou & Grigorieff, 2015). By subclassing the `AbstractCTF`, custom implementations may be created downstream to the core cryoJAX codebase. This is demonstrated in Code example 1. By implementing a method for the aberration function, this custom CTF may be used with the rest of cryoJAX to simulate images. Though this is a simple example, the principle applies generally. For example, it is possible to subclass the `AbstractVolumeRepresentation` and create custom volume representations. This makes cryoJAX a flexible modeling language for cryo-EM image formation and a powerful tool for cryo-EM research.

Using cryoJAX in practice revolves around building an image formation model. This process is demonstrated by the code presented in Figure 2 B. First, the `GaussianMixtureVolume` class is instantiated in three steps: use `cryojax.io` to load atom positions and atomic numbers, call the helper class `PengScatteringFactorParameters` to load tabulated electron scattering factor parameters, and instantiate the `GaussianMixtureVolume`. Next, an `EulerAnglePose` is instantiated. This is implemented in a standard cryo-EM convention: the first euler rotation *ϕ* is about the z-axis, then *θ* about the y-axis, and finally *ψ* about the z-axis. Finally, the `AstigmaticCTF` is selected and wrapped into a `ContrastTransferTheory` class, which determines how to apply the CTF to the projection.

Finally, the image formation model is configured, instantiated, and simulated. Image formation models, like any class in cryoJAX, follow a similar theme described in Table 1. The abstract interface that performs simulation is called the `AbstractImageModel`, and `make_image_model` is a utility function for instantiating core implementations in cryoJAX.

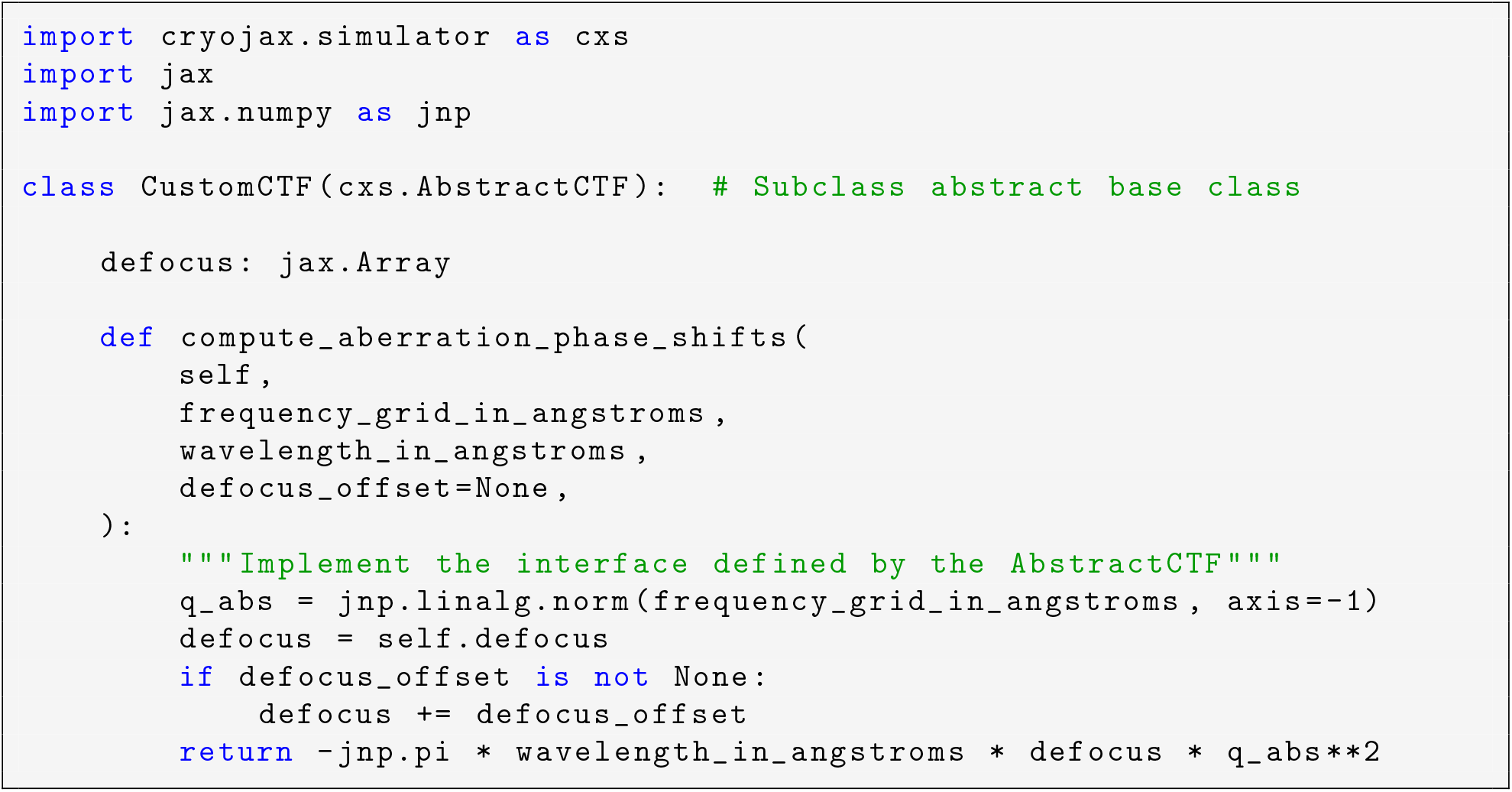

Code example 1: Creating a custom implementation of a CTF. The `AbstractCTF` interface only has one method to implement for the frequency-dependent aberration phase shifts, *χ*. We demonstrate implementing the defocus term of a CTF: 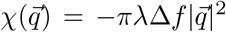, where Δ*f* is defocus, λ is the wavelength of incident electrons, and 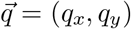 is an in-plane spatial frequency vector. This class can be used to simulate images as if it were included in the core cryoJAX library. The same principle applies to other interfaces in cryoJAX, such as the `AbstractVolumeRepresentation`.

By default, this is the `LinearImageModel`, which is an implementation of Equation 1 without scaling to physical units using σ_e_. An implementation for simulating images with no CTF applied is also included, as well as implementations for simulating physical quantities using Equation 1 (i.e. contrast, intensity and electron counts). We have demonstrated the use of these tools for the simulation of single-particle images. While micrographs can also be simulated by requesting a large image, it would be straightforward to implement a micrograph or movie frame simulator that includes effects such as defocus gradients and ice movement (Himes & Grigorieff, 2021).

To demonstrate that the model in Figure 2 B can describe the data, we compare a simulated image against an image from EMPIAR-10833, an experimental dataset of thyroglobulin in Figure 2 C. We display radially averaged power spectra for each image, which show Thon ring overlap at low frequencies. We calculated a cosine similarity of 0.97 to quantify the overlap, which indicates a high degree of similarity between the experimental and simulated power spectra. To generate the simulated image, we perform cryoSPARC consensus reconstruction to estimate pose and CTF parameters (Appendix B.1) and load them using the cryoJAX STAR file reader. Using PDB entry 6SCJ, we simulate the image using the `GaussianMixtureVolume` class and add colored noise with a power spectrum described in Appendix B.2. We further demonstrate the interoperability of cryoJAX with standard cryo-EM software in Appendix C, where we use cryoJAX to simulate synthetic thyroglobulin images and use the resulting metadata in the form of a STAR file to reconstruct the molecule using RELION and cryoSPARC.

Image simulation features in cryoJAX are not limited to implementations based on Equation 1. For example, cryoJAX has preliminary support for simulation of contrast from large specimens via Ewald sphere extraction (Wolf *et al*., 2006) and explicit wavefunction calculation for strong phase imaging (Vulović *et al*., 2014; Rickgauer *et al*., 2017). CryoJAX is flexible enough to support more advanced features for simulating accurate contrast, such as dose-dependence (Grant & Grigorieff, 2015), solvent effects (Vulović *et al*., 2013; Himes & Grigorieff, 2021), more accurate electrostatic potentials (Vulović *et al*., 2013; Shtyrov *et al*., 2025), and multislice wave propagation (Hall *et al*., 2011; Vulović *et al*., 2013; Himes & Grigorieff, 2021; Parkhurst *et al*., 2021). CryoJAX also implements interfaces for the simulation of Gaussian white noise and colored noise from arbitrary power spectra. This can be generalized in the future to include other distributions and physical models of structured noise (Parkhurst *et al*., 2024). More accurate models of high-resolution information in cryo-EM images will improve existing data analysis procedures and enable new applications.

### 2.3 Using cryoJAX for data analysis

With cryoJAX as a modeling framework for cryo-EM data, the scientific computing tools of JAX may be leveraged for building data analysis applications. JAX is a popular array-oriented numerical computing library in Python for *function transformations*: operations that act on functions to return new functions. Three core transformations power JAX applications: just-in-time (JIT) compilation, automatic vectorization, and automatic differentiation.

These transformations make JAX a powerful and flexible framework for building computationally efficient data analysis applications. JIT compilation in JAX gives code the flexibility of Python with the performance of compiled languages like C++ and CUDA. Automatic vectorization is capable of transforming highly non-trivial functions, even entire programs, to their vectorized versions. Automatic differentiation makes it possible to formulate gradient-based optimization of a loss function for staggering numbers of parameters. Finally, the JAX JIT compiler can automatically determine how to efficiently distribute entire workflows across GPUs. In cryoJAX, there are no assumptions of how users will use these function transformations; cryoJAX is just an image simulator, yet these transformations make it possible to build highly complex programs.

Alongside JAX, CryoJAX is built on the library Equinox (Kidger & Garcia, 2021). Equinox is a popular JAX library for PyTorch-like classes that smoothly integrate with JAX function transformations. In cryoJAX, these classes are used to build the abstract base class framework for modular image simulation. Generically, they are used to represent functions whose parameters interact with JAX transformations. Equinox provides an elegant framework for using cryoJAX classes with JAX transformations.

Using Equinox, image simulation is easily JIT compiled (Code example 2). To do so, first build the relevant cryoJAX modeling components (i.e. the volume, CTF, and pose). This process is demonstrated in Figure 2 B. Then, define a Python function that accepts these modeling components and returns an image. By applying Equinox’s Python decorator `equinox.filter_jit` (a lightweight wrapper around `jax.jit`), this function is transformed into a JIT compiled version. This function may be called in the same way as in the original function signature to simulate an image.

To demonstrate JIT compilation, we simulated 200 by 200 images at 2 Å pixel size in cryoJAX using the thyroglobulin structure used in Figure 2 C. For simplicity, only the C-alpha atoms are used. Images are simulated using an implementation of Eq 1: the rotation **R** is implemented using the `EulerAnglePose` class, the CTF sin *χ* is implemented using the `AberratedCTF` class, and we test both voxel (i.e. `FourierVoxelGridVolume`) and atom (i.e. `GaussianMixtureVolume`) volume representations U. On a NVIDIA A100 GPU (A100-SXM4-40GB), using cryo-JAX’s `FourierVoxelGridVolume` class for image simulation took approximately 367.3 ± 1.5 ms without JIT compilation and 0.45 ± 0.07 ms with JIT compilation, approximately an 800 times speed-up. CryoJAX’s `GaussianMixtureVolume` class using the 5,382 C-alpha atoms took 253.5±1.4 ms without JIT and 0.58 ± 0.09 ms with JIT, approximately a 435 times speed-up. In Appendix D, we summarize this test and include an analogous test on a CPU. Note that JAX is not optimized for cases without JIT compilation, so these speed-ups should not be compared with other Python GPU scientific computing frameworks without JIT.

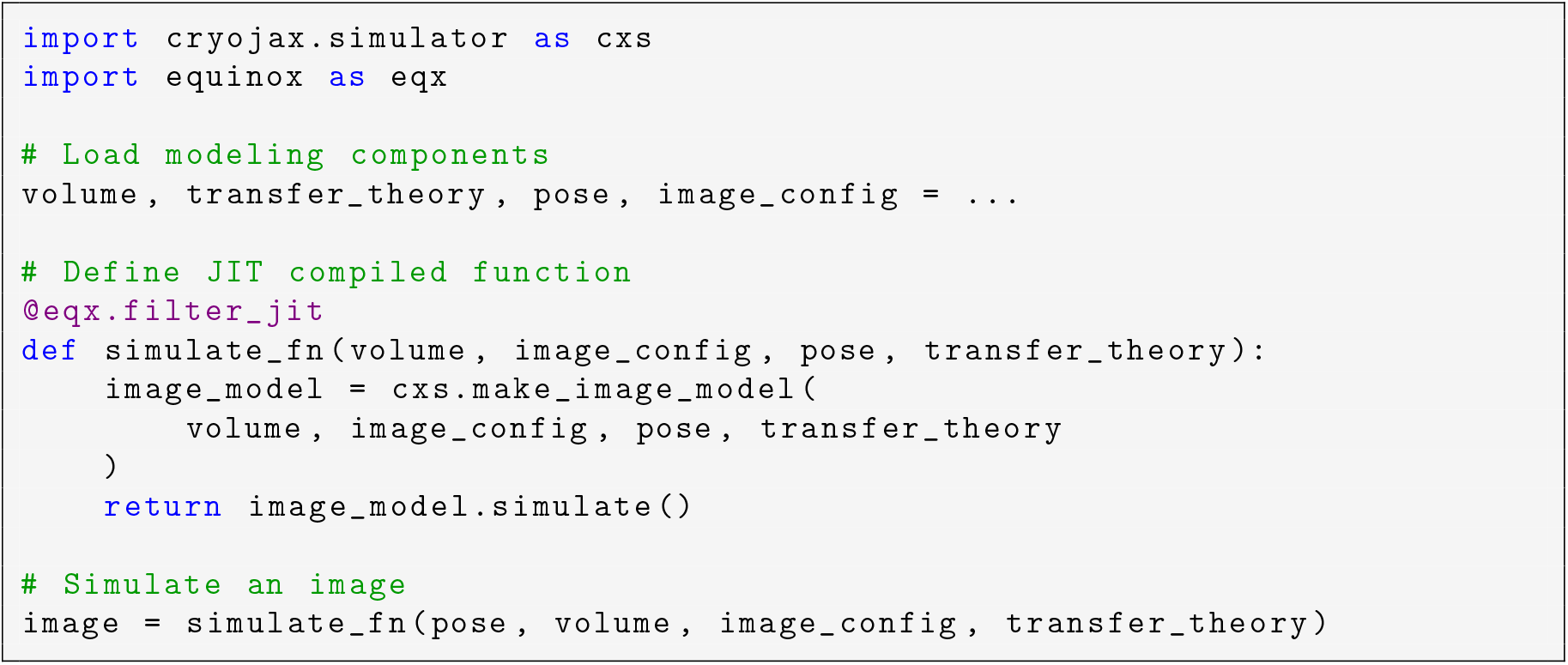

Code example 2: Simulation of an image using Equinox, a library for integrating JAX with Python classes, for just-in-time (JIT) compilation. JIT compilation is a key feature of the JAX library and is used to dramatically improve program performance.

Given modeling components whose parameters are arrays with a batch dimension, simulation of a single image in Code example 2 can be easily transformed to a vectorized version using `equinox.filter_vmap`. We test a JIT compiled and vectorized image simulation function over the pose on the same GPU hardware as for the single image case. Again using cryoJAX’s `FourierVoxelGridVolume` class for Fourier slice extraction, simulation of 100 images took approximately 2.6 ± 0.1 ms, only an approximate 5 times slowdown from the single image case. For the `GaussianMixtureVolume` case it took 12.5 ± 0.2 ms, only an approximate 20 times slowdown. These performance gains should decrease with larger image number due to GPU memory saturation. To investigate this phenomenon and further quantify performance, we extend this test in Appendix E for the `FourierVoxelGridVolume` case and conduct sweeps over box size and the number of images simulated.

It is also straightforward to define a loss function with respect to a cryo-EM image and use a JAX automatic differentiation function transformation to compute gradients (Code example 3). Consider the task of computing the cross-correlation score between simulated and real images and taking gradients with respect to the protein atom positions. First, the user loads a cryo-EM image in Python and again builds relevant modeling components. Given atom positions, a Python function is defined for the cross-correlation loss. This is done in three steps: finish building relevant modeling components (i.e. the volume), create the image formation model and simulate an image, and compute the cross-correlation score. The JAX transformation is implemented through the Python decorator `equinox.filter_value_and_grad` (a lightweight wrapper for `jax.value_and_grad`) to transform the loss function to one that returns its value and gradients with respect to atom positions.

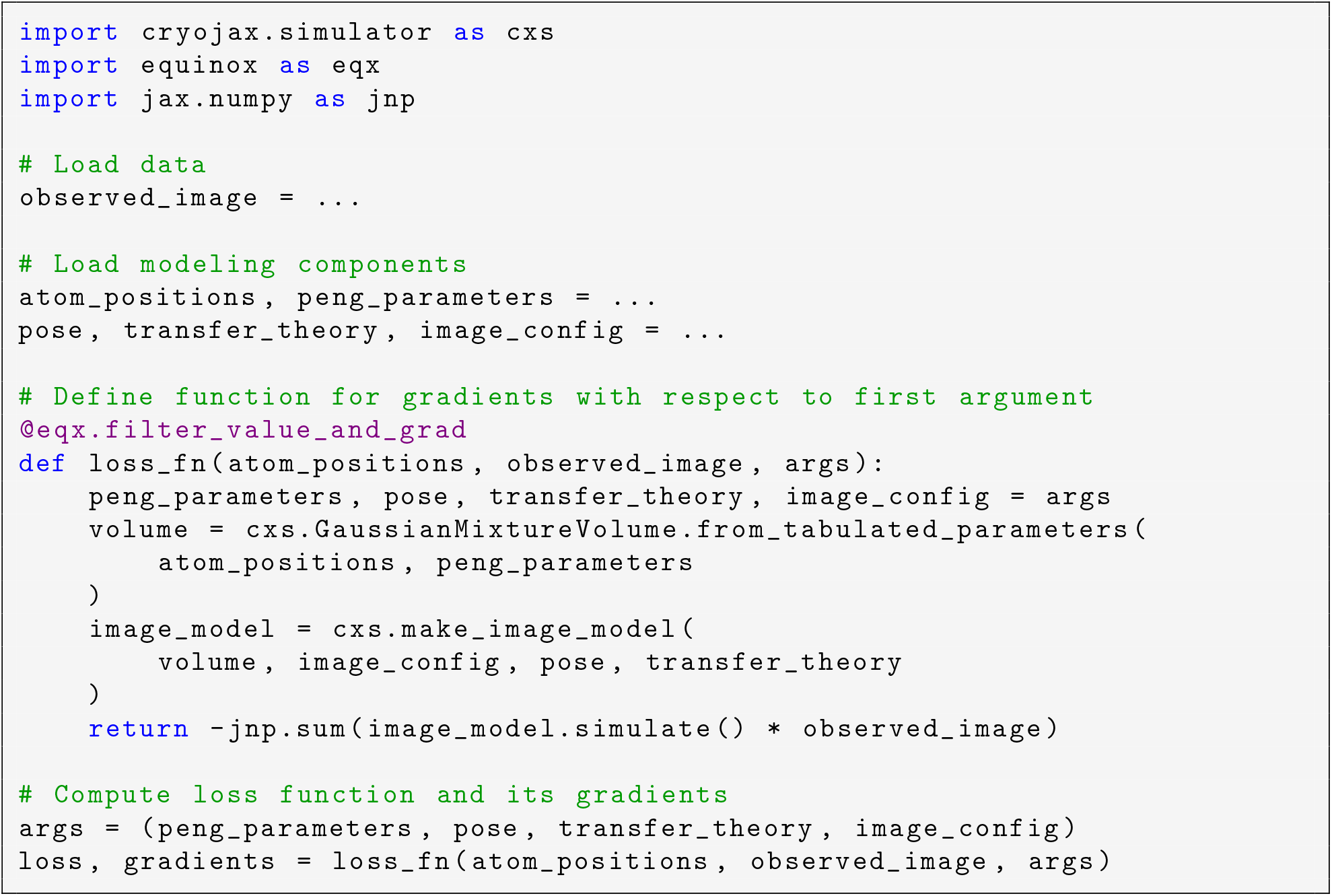

Code example 3: Compute gradients using automatic differentiation with respect to atom positions of a cross-correlation loss function. Despite that atom positions can be in the hundreds of thousands, automatic differentiation makes it possible to formulate gradient-based optimization for this loss function.

With numerical differentiation using finite differences, it would be prohibitively computationally expensive and not numerically stable to compute gradients with respect to atom positions, which can be up to hundreds of thousands of parameters. Automatic differentiation computes exact numerical gradients with computational cost that scales with the function evaluation, rather than with the number of parameters differentiated. This technology has enabled the revolution of machine learning (LeCun *et al*., 2015), where neural networks may have millions of parameters. Combined with the flexibility of cryoJAX, JAX automatic differentiation can be used to formulate gradient-based optimization of many parameters, such as those parametrizing a protein’s structure.

#### Example: structure refinement using automatic differentiation

To demonstrate the utility of JAX automatic differentiation for cryo-EM data analysis, we formulate an optimization procedure that refines protein structure at known pose and CTF parameters (Figure 3). Using cryoJAX we generate a synthetic dataset of 100 images at SNR = 0.1 using a bent conformation of thyroglobulin obtained from MD simulation (Astore *et al*., 2025) (Figure 3 A). To simulate the images, the volume is represented as a mixture of Gaussians centered on the thyroglobulin C-alpha atom positions with equal weight and variance.

**Figure 3.**
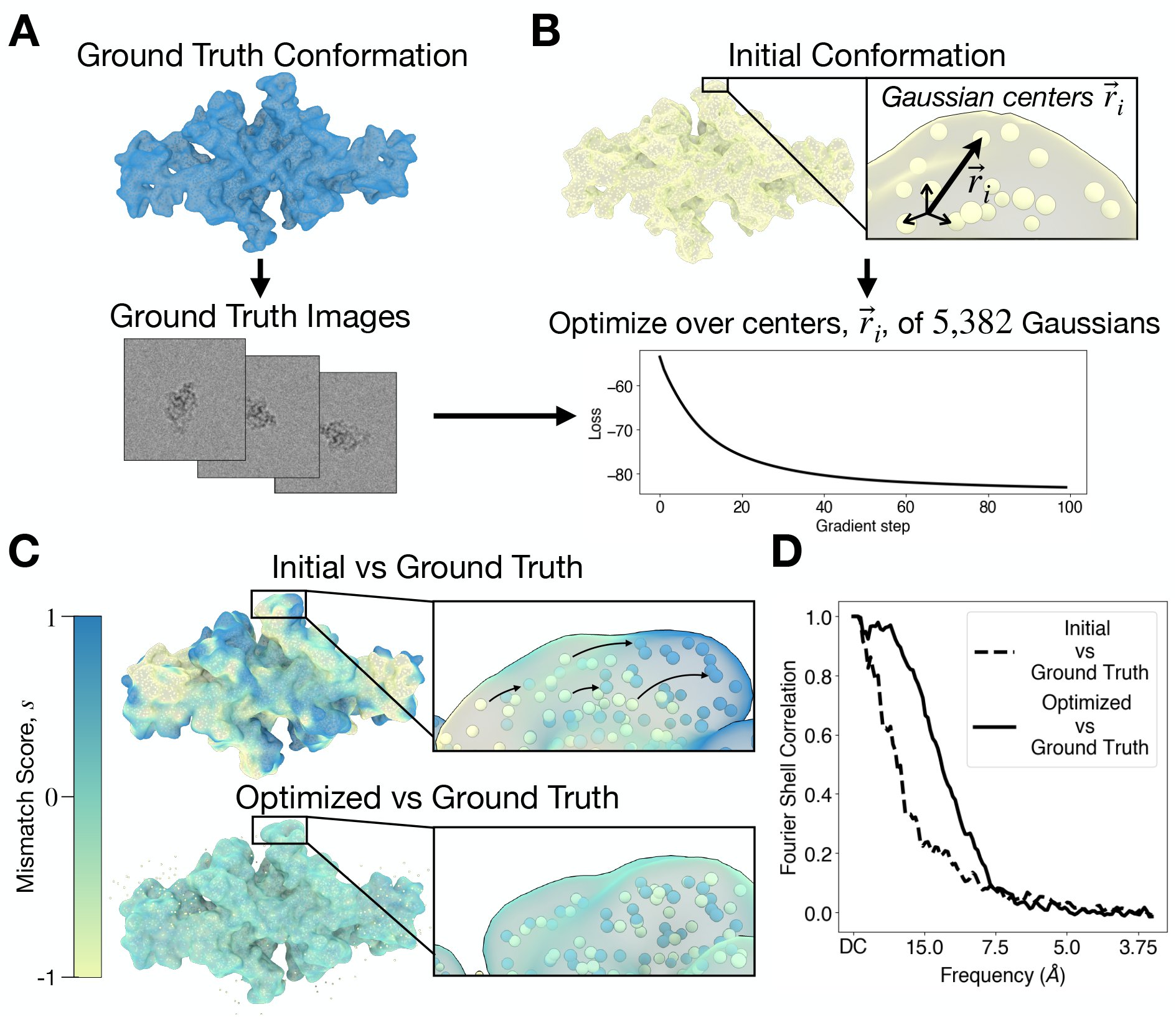
An application of automatic differentiation. **A.** A structure of thyroglobulin is used to generate a stack of synthetic images used as ground truth. Images are simulated by representing C-alpha atoms as Gaussians with equal variance. The centers of the Gaussians are represented by spheres while the volumes these Gaussians create are shown as a transparent isosurface. **B**. Starting from a structure of thyroglobulin deformed from its original conformation, the centers of the Gaussians are optimized using gradient descent of a loss function. **C**. After optimization, the Gaussians capture the low-resolution structure of the ground truth. This is illustrated by computing the sum of the volume with its ground truth before and after optimization and rendering it as an isosurface colored by the mismatch score, *s*. *s* ≈ 0 indicates little mismatch, while *s* ≈ 1 or *s* ≈ −1 indicate significant mismatch. **D**. The Fourier shell correlation captures the improved agreement at low-resolutions between the optimized structure and the ground truth.

Starting from a straightened conformation of thyroglobulin, we perform gradient-based optimization over the 5, 382 Gaussian centers to recover the ground truth conformation (Figure 3 B). To perform the optimization, we use a cross-correlation loss function at fixed CTF and pose parameters vectorized over all 100 synthetic images and parameters using the JAX transformation `jax.vmap`. We optimize atom positions using the AdaBelief algorithm (Zhuang *et al*., 2020) in the JAX optimization library Optax (DeepMind *et al*., 2020). After 100 gradient steps with respect to 5, 382 Gaussian centers, the loss function reaches a minimum. For images of size 200 by 200, this runs in less than 5 minutes on a laptop CPU.

This optimization procedure successfully captures the low resolution structure of the ground truth conformation. Figure 3 C illustrates the mismatch between the ground truth and both the initial and the optimized conformations. We display isosurfaces rendered with ChimeraX of the sum of the ground truth and the initial (upper panel) or optimized (lower panel) volumes, which are discretized on a 3D real-space voxel grid using cryoJAX. We color these isosurfaces by what we call the mismatch score, *s* = 2 *U*_0_/(*U*_0_ + *U*) − 1 where *U*_0_ is the ground truth volume and *U* is either the initial or optimized volume. The mismatch score for the initial conformation has distinct regions where *s* ≈ 1 or *s* ≈ −1, indicating significant mismatch. For the optimized conformation, the mismatch score is dominated by *s* ≈ 0. Therefore, the optimized and ground truth structures have little mismatch. This is further captured by the Fourier shell correlation (FSC) in Figure 3 D. The FSC between the ground truth and the optimized volume decorrelates at higher frequencies than the FSC between the ground truth and initial structure.

This example demonstrates the flexibility of cryoJAX and the power of JAX for cryo-EM data analysis. However, it also demonstrates the flexibility of JAX. CryoJAX is simply an image simulator, yet it is possible to build complex and broad data analysis applications using `jax.jit`, `jax.vmap`, and `jax.grad`. In other words, the cryoJAX library code can remain simple because JAX transformations make it possible for application code to specify how cryoJAX will be used in practice. With cryoJAX, JAX, and Equinox, cryo-EM data analysis for broad scientific application may be prototyped and deployed at scale.

## 3 Discussion

Cryo-EM is a rapidly growing technique for the biophysical sciences. As it is used for increasingly broad scientific goals, there is a need for software designed for building and testing new ideas that may be deployed for studying experimental data at scale. We argue that cryoJAX fulfills this role as a flexible image simulation library that integrates with JAX.

In this article, we motivate the choice of JAX on the basis of its computational capabilities. However, PyTorch is another Python framework with a larger user-base and similar capabilities compared to JAX. Our justification of JAX over PyTorch is largely because the flexibility of the JAX programming model enables the specific aims of the cryoJAX project. Particularly when JAX is combined with Equinox, JAX enables cryoJAX’s design goals as a library that is flexible, intuitive, and straightforward to maintain. Although PyTorch has recently implemented JAX-like transformations via `torch.func`, its ability to handle transformations in complex settings lags behind those of JAX at the time of writing. For users seeking to leverage the vast ecosystem of PyTorch software, a growing number of packages exist to interoperate between the two frameworks (Qi *et al*., 2025; Normandin, 2026).

Comparing cryoJAX to other standalone simulation packages is challenging, as these typically differ from cryoJAX in their primary focus. At the time of publication, the closest analog to cryoJAX in the literature is TeamTomo (Burt & Giammar, 2026). Like cryoJAX, TeamTomo is a Python open source project actively being used for the development of new cryo-EM data analysis methods (Burt *et al*., 2024; Sanchez-Garcia *et al*., 2025; Giammar *et al*., 2025; Shah *et al*., 2025). While there is some overlap in specific functionality, cryoJAX is intended as a framework for the development of cryo-EM methods for emerging scientific applications, while TeamTomo focuses on in situ cryo-EM/ET method development and simplifying general purpose cryo-EM/ET scripting. TeamTomo and cryoJAX implement image processing functionality using PyTorch and JAX respectively to provide GPU accelerated computation and automatic differentiation. CryoJAX is distributed as a single package with a layered API focused on differentiable image simulation as a framework for downstream data analysis. TeamTomo instead distributes a number of small libraries which implement file IO operations, low level image analysis primitives, and higher level image processing algorithms. The two efforts complement each other and both show great promise as engines for innovation in the field.

While the primary aim of cryoJAX is to facilitate the development of cryo-EM data analysis, the challenge of using cryo-EM for new scientific applications poses additional challenges. Modeling cryo-EM images themselves is an open problem, and recently other simulation libraries have made advancements in detailed modeling of the image formation process (Himes & Grigorieff, 2021; Parkhurst *et al*., 2021). However, evaluating the utility of these advancements has been inhibited by the challenge of deploying these simulators at scale and implementing controls against standard methods. These studies can be implemented within cryoJAX, whose modularity and integration with JAX enables studies of image formation itself. By leveraging JAX’s tools for statistical inference and machine learning, it is also possible to use data-driven approaches to design new simulation methods, such as for the simulation of noise distributions *in situ*. Improved agreement between simulated and real images will improve existing data analysis and possibly enable new techniques that leverage information at high-resolutions.

The flexibility and accessibility of cryoJAX has led to its adoption in several projects during its development. A simple use case has been to create synthetic datasets (Astore *et al*., 2024), but it has also been used to refine molecular ensembles from experimental data (Silva-Sánchez *et al*., 2024) and explore the use of optimal transport loss functions for cryo-EM (Woollard *et al*., 2025*a*).

The cryoJAX codebase is available on Github at https://github.com/michael-0brien/cryojax under the GNU Lesser General Public License. It is uploaded to the Python Package Index (PyPI) and is installable using pip. The documentation and testing suite are currently in development. CryoJAX is a community-led open source project; anyone is welcome to contribute.

## Acknowledgements

We would like to thank Nikolaus Grigorieff for insight and guidance on algorithms for cryo-EM image simulation throughout this project. M.J.O. would like to thank Patrick Kidger, Louis Desdoigts, and other members of the JAX open source scientific software community for helpful discussions that have informed the design of the cryoJAX codebase. M.J.O would also like to thank Alister Burt for assisting with the comparison between TeamTomo and cryoJAX. Finally, we would like to thank Luke Evans, Wai-Shing Tang, Shuyu van Kerkwijk, Minhuan Li, Anupam Anand Ojha, and all those who contributed to the codebase and gave feedback during the library’s development.

## Contributions

M.J.O. conceived of the study, developed the codebase, and wrote the manuscript. D.S.S. developed the codebase and contributed to the manuscript. G.W. contributed to the codebase and the manuscript. K.J. contributed to the manuscript. P.C., S.H., and D.J.N. provided supervision. E.H.T. provided supervision and contributed to the codebase. M.A.A. provided supervision and wrote the manuscript.

## Data availability

The cryoJAX codebase and documentation can be accessed from the following link: https://github.com/michael-0brien/cryojax.

## A Computing the electrostatic potential from an atomic model

A scattering calculation for cryo-EM image formation starts with the electrostatic potential energy distribution of the scatterer, denoted 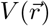. The image simulated in Figure 2 uses a cryoJAX core implementation for computing electrostatic potential energy distributions. In this appendix, we outline this model.

CryoJAX uses a convention of the following re-scaling of the potential energy into dimensions of length^−2^:

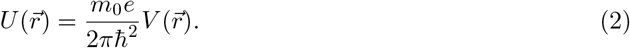

for the electron rest mass *m*_0_, electron charge *e*, and planck’s constant ħ. This gives the following Schrodinger equation,

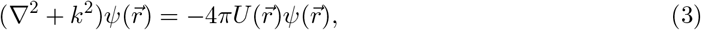

where *k* is the wavenumber of the incident electron beam.

In order to compute the potential energy distribution of a single atom, we assume that the potential is weak relative the energy of the incident beam, i.e. *U* << *k*^2^. Further, we only consider single scattering events about individual atoms. This gives the first Born approximation, where the potential can be computed as the inverse Fourier transform of an electron scattering factor (the scattering amplitude of an electron by an atom). We will denote an electron scattering factor as 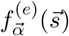, where 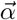 is a vector that parametrizes the scattering factor and 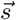 is referred to as the scattering vector. Then the potential can be computed from the first Born approximation as

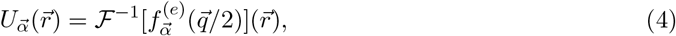

where the Fourier convention is defined 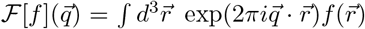 for the 3D wave vector coordinate 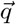. Note that 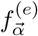 has dimensions of length and the scattering vector is related to the wave vector as 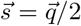

Many works have parametrized and tabulated electron scattering factors across individual atom types. In cryo-EM, electrostatic potentials for a protein are approximated as the sum of electrostatic potentials computed from tabulations for each atom. While cryoJAX can be extended to handle alternate scattering factor parametrizations (or even alternate approximations of the potential), cryoJAX currently includes a core concerete implementation for the approximation from Peng et al. (1996) (Peng *et al*., 1996). This computes an electron scattering factor as a sum of 5 Gaussians. The parameters for each Gaussian include amplitudes 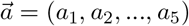 and variances 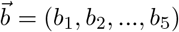. Also defining an additional empirical B factor, the expression for the scattering factor is

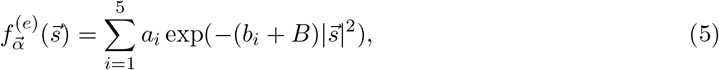

where 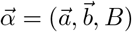 is the vector of parameters.

Substituting these scattering factors into Equation 4 gives the following potential energy distribution centered at the origin,

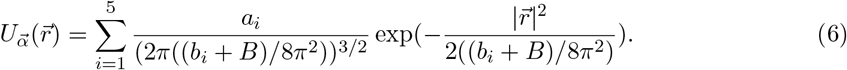

Since the variance of the potential from a single atom is typically on the order of the camera pixel size or less (i.e. the Gaussians are short ranged), discretization effects often cannot be neglected and Eq. 6 must be averaged within individual voxels. With voxel size Δ*r* and voxels indexed by *ℓ*, this average is expressed as,

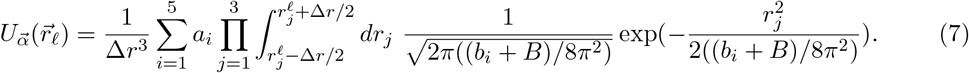

In practice, this is computed analytically with error function evaluations,

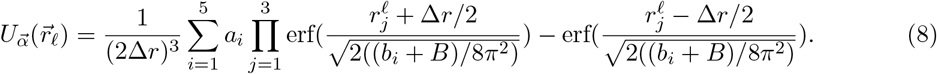

Finally, to compute the potential energy distribution of the scatterer, we treat all atoms independently. Therefore, a collection of M independent atoms parametrized by vectors 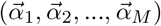 is a superposition of individual atom potentials at positions 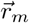

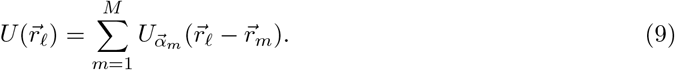

## B Methods: Processing of experimental and simulated images of thyroglobulin

### B.1 Processing EMPIAR-10833

Experimentally collected movies deposited on EMPIAR (EMPIAR-10833) (Kim *et al*., 2021) were gain-corrected and processed with MotionCorr2 (Zheng *et al*., 2017). We picked particles using a blob picker with radius of 180-200 Å, and these coordinates were used to extract particles with a box size of 448 pixels. Two rounds of 2D classification were used to remove junk particles. The cleaned particles were subjected to homogeneous refinement and local refinement with C2 symmetry for a reconstruction with a resolution of 3.0 Å. All processing of EMPIAR-10833 data was performed in cryoSPARC (Punjani *et al*., 2017*b*).

### B.2 Simulating synthetic images

The simulated image in Figure 2 C is generated from pose and CTF parameters drawn from the reconstruction procedure described in Appendix B.1 with PDB entry 6SCJ. The colored noise model used to simulate the image has a two-parameter power spectrum

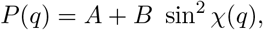

where 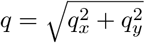 is the in-plane spatial frequency vector magnitude and sin *χ*(*q*) is the CTF. This model for the power states that each Fourier mode (*q*_*x*_, *q*_*y*_) of the noise is an independent Gaussian random variable *X*(*q*_*x*_, *q*_*y*_) = *X*_*A*_ + *X*_*B*_(*q*_*x*_, *q*_*y*_), where 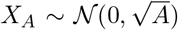 has constant variance and 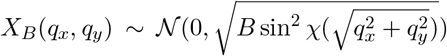 has variance dependent on the CTF. We use a model with CTF dependence because the solvent background in cryo-EM images is CTF dependent. We empirically choose the noise model parameters *A* and *B* and the standard deviation of the signal so that the power spectra of the simulated image and experimental image are well-aligned.

## C Reconstructing synthetic images in RELION and cryoSPARC

To demonstrate the interoperability of the cryoJAX code with typical cryo-EM analysis pipelines, we performed a reconstruction of a synthetic single-particle dataset with both RELION and cryoSPARC. Using a PDB structure of the thyroglobulin molecule, we created a synthetic dataset of 100 000 images using a uniformly random pose distribution with a signal to noise ratio of 0.01 (Figure 4 A). The associated STAR file with defocus values and particle poses was used to perform fourier-slice insertion with the *relion-reconstruct* command and the *“Homogeneous Reconstruction Only”* job in RELION and cryoSPARC respectively (Figure 4 B). For both cases this resulted in the correct reconstruction of the input thyroglobulin molecule, demonstrating the compatibility of the simulation methods and conventions used in cryoJAX and those in established cryo-EM software packages (Figure 4 C).

**Figure 4.**
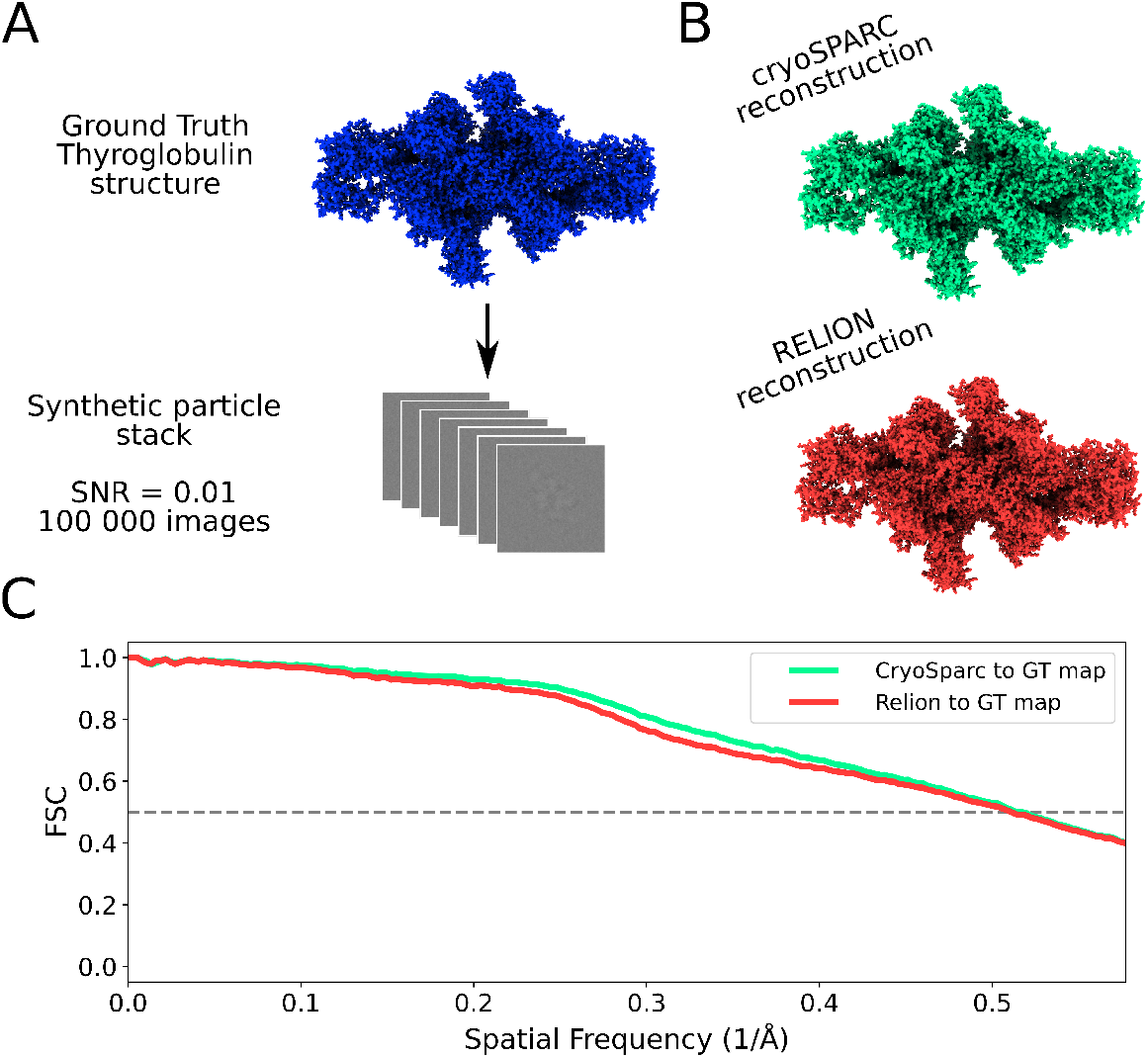
The faithful reconstruction of a cryoJAX generated particle stack by RELION and cryoSPARC. **A.** A PDB structure of thyroglobulin was used to generate 100 000 single-particle images. **B**. The images and their associated metadata in a STAR file were read by RELION and cryoSPARC. This information was used to accurately reconstruct a map of thyroglobulin. **C**. The RELION and cryoSPARC reconstructed voxel maps had a fourier shell correlation to the ground truth map of 1.96 Å and 1.95 Å respectively. This example shows the correspondence and interoperability between the conventions of cryoJAX and popular cryo-EM image processing software.

## D Demonstrating JAX Just-In-Time (JIT) compilation

In Section 2.3, we demonstrated cryoJAX image simulation with and without JAX just-in-time (JIT) compilation. We simulated a 200 × 200 image using the 5, 382 C-alpha atoms of thyroglobulin used in Figure 2 C. Timings are shown for JAX configured to use both a 32-core CPU (AMD Ryzen Threadripper 3970X 32-Core) and a NVIDIA GPU (A100-SXM4-40GB), and we test implementations where projections are obtained from extracting Fourier slices (i.e. `FourierVoxelGridVolume`) and analytically integrating Gaussians on the plane (i.e. `GaussianMixtureVolume`). Timings are averaged over 100 trials, and floating point calculations are performed in 32-bit precision. Memory transfers between GPU and CPU and are not included in compute times.

## E Benchmarking vectorized image simulation

In Section 2.3, we present a test case that computes batches of images by converting an image simulation function to its vectorized version using `jax.vmap`. Figure 5 expands on this test for the case of cryoJAX’s implementation of Equation 1 using Fourier slice extraction. We compute benchmarks for vectorized image simulation from 1 − 10^4^ images and successive box size doublings from 64 − 512 pixels per dimension. Simulation is performed on an NVIDIA GPU (H100-80GB), and compute times are averaged over 100 function evaluations. The image simulation implementation includes Fourier slice extraction via linear interpolation, CTF and translational phase shift application, and inverse Fourier transform. The time to render the volume as a voxel map in the Fourier domain is not included, nor is GPU to CPU memory transfer. In practice, true compute times may depend strongly on the implementation in a given data analysis application.

**Figure 5.**
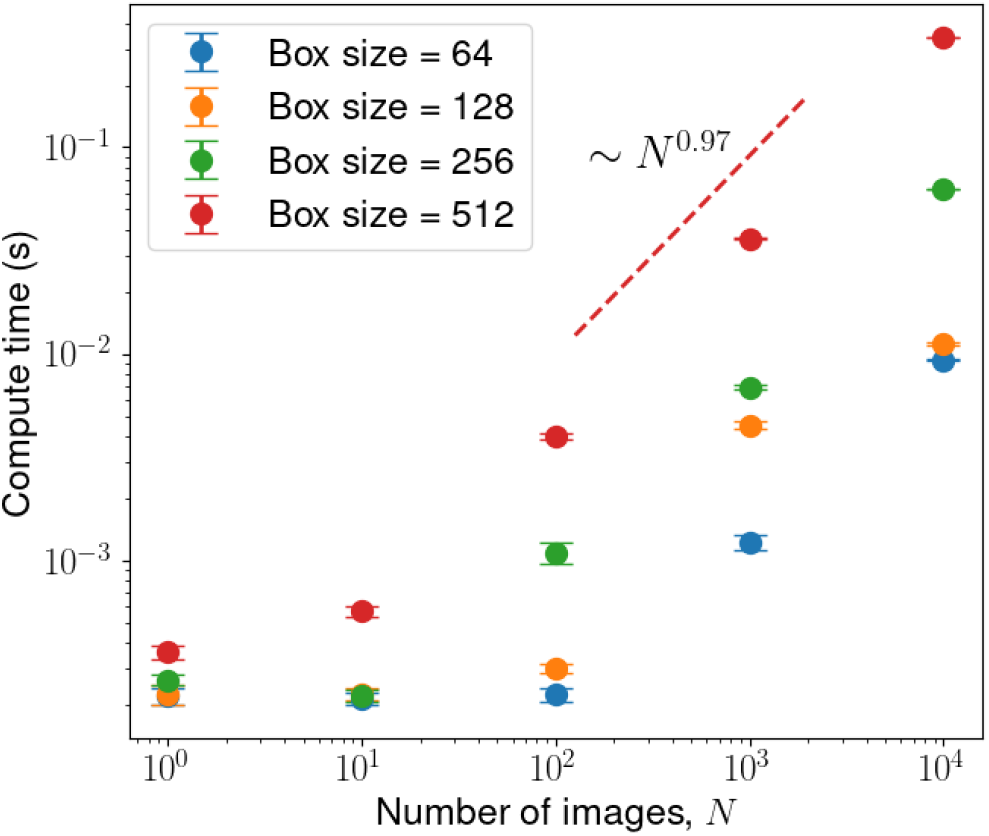
Scaling of vectorized image simulation over rotations via Fourier slice extraction. Compute times of cryoJAX vectorized image simulation using Fourier slice extraction demonstrate the performance capabilities of cryoJAX. Simulating at least 10 images took under a millisecond for all box sizes, and simulating 10^4^ images remained under 100 milliseconds for all but the box size equal to 512 case. In all cases, a power law emerges after a critical number of images. For the box size 512 case, this power law shows near-linear scaling, suggesting that performance gains have diminished due to GPU memory saturation. The 80 gigabytes of available GPU memory is exhausted after 10^4^ images, so it becomes necessary to implement additional strategies such as iteration or deployment across multiple GPUs. These calculations were performed on an H100-80GB NVIDIA GPU with a 64-core Icelake Intel CPU.

